# High-Resolution Satellite Imagery to Assess *Sargassum* Inundation Impacts to Coastal Areas

**DOI:** 10.1101/2020.08.10.244004

**Authors:** William J. Hernandez, Julio M. Morell, Roy A. Armstrong

## Abstract

A change detection analysis utilizing Very High-resolution (VHR) satellite imagery was performed to evaluate the changes in benthic composition and coastal vegetation in La Parguera, southwestern Puerto Rico, attributable to the increased influx of pelagic *Sargassum spp* and its accumulations in cays, bays, inlets and near-shore environments. Satellite imagery was co-registered, corrected for atmospheric effects, and masked for water and land. A Normalized Difference Vegetation Index (NDVI) and an unsupervised classification scheme were applied to the imagery to evaluate the changes in coastal vegetation and benthic composition. These products were used to calculate the differences from 2010 baseline imagery, to potential hurricane impacts (2018 image), and potential *Sargassum* impacts (2020 image). Results show a negative trend in Normalized Difference Vegetation Index (NDVI) from 2010 to 2020 for the total pixel area of 24%, or 546,446 m^2^. These changes were also observed in true color images from 2010 to 2020. Changes in the NDVI negative values from 2018 to 2020 were higher, especially for the Isla Cueva site (97%) and were consistent with the field observations and drone surveys conducted since 2018 in the area. The major changes from 2018 and 2020 occurred mainly in unconsolidated sediments (*e.g.* sand, mud) and submerged aquatic vegetation (*e.g.* seagrass, algae), which can have similar spectra limiting the differentiation from multi-spectral imagery. Areas prone to *Sargassum* accumulation were identified using a combination of 2018 and 2020 true color VHR imagery and drone observations. This approach provides a quantifiable method to evaluate *Sargassum* impacts to the coastal vegetation and benthic composition using change detection of VHR images, and to separate these effects from other extreme events.

## Introduction

The floating brown algae *Sargassum* sp. provide an essential marine habitat in the open ocean. However, since 2011, there has been a dramatic increase in *Sargassum* biomass in the tropical Atlantic Ocean and Caribbean Sea and consequently, massive accumulations of *Sargassum* have been reported along the Eastern and Western Caribbean and Florida coasts [1, 2]. From mid-2014 until the end of 2015, the Mexican Caribbean coast experienced a massive influx of drifting *Sargassum* that accumulated on the shores [3]. This build-up of decaying beach-cast material and nearshore accumulations produced murky brown waters that were called *Sargassum*-brown-tides (Sbt) with persistent reduction in light, oxygen (hypoxia or anoxia) and pH. In addition to the water quality impacts, they found that seagrass meadows dominated by *Thalassia testudinum* were replaced by calcareous rhizophytic algae and drifting algae and/or epiphytes, which resulted in a 61.6–99.5% loss of below-ground biomass and reduced value to coastal blue carbon storage. High levels of eutrophication were still present after one year of the Sbt and near-shore corals suffered total or partial mortality.

Remote sensing have been employed by many researchers to map general benthic habitat types (*e.g.*, sand, seagrass, coral reefs, hard substrate) in coral reef environments [4,5,6] and coastal vegetation [7,8] in Puerto Rico. Also, coral reef habitat maps, based on remotely sensed data, are a fundamental tool for management because they summarize ecologically meaningful information across extensive geographic scales in a cost-effective manner [9]. In addition, changes to the vegetation coverage from extreme events (*e.g.* hurricanes) have been documented using remotely sensed data for the estimation of NDVI [7,8].

Initial observations in La Parguera area (SW Puerto Rico) suggest that *Sargassum* influx and accumulations have increased since 2011, with a significant increase in the peak years of *Sargassum* influx to the Caribbean [1]. Additionally, original benthic cover and composition, as well as coastal vegetation may have been displaced and/or significantly impacted by persistent *Sargassum* accumulations in near-shore areas. The purpose of this study was to evaluate the changes in benthic composition and coastal vegetation in La Parguera area due to *Sargassum* influx and accumulations in cays, bays, inlets and near-shore environments.

### Study Area

The study area in La Parguera was selected due to the observed increased frequency of events resulting in the accumulation of *Sargassum* at some of the coral reefs and mangrove-fringed coasts. The constraints to hydrodynamic activity, provided by reefs and mangrove keys in the study area, coupled to the prevalence of south-easterly winds makes the study area particularly vulnerable to Sargasso accumulation The benthic and coastal vegetation area includes the Boquerón State Forest and La Parguera Natural Reserve, managed by the Puerto Rico Department of Natural and Environmental Resources (DNER) (Figure 2). The area of La Parguera is recognized for the exceptional value of its marine resources, which includes extensive coral reef ecosystems, seagrass beds, coastal mangrove fringe and mangrove islands, and two bioluminescent bays [10]. The study areas were further subdivided into the Isla Cueva (IC), Isla Guayacán (IG), and Boquerón State Forest (BF) areas (Figure 1).

**Figure 1:**
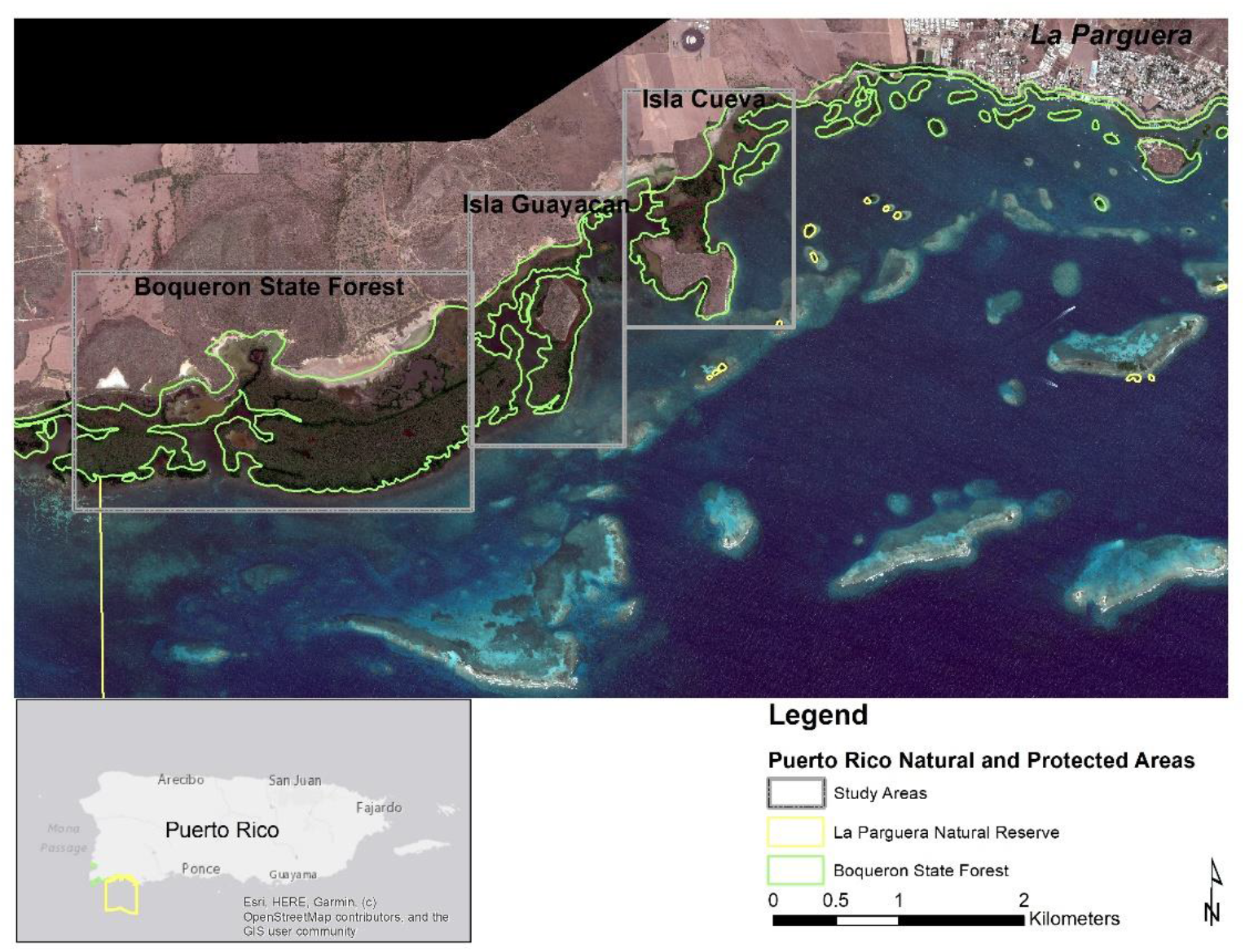
Map showing the study areas for La Parguera. The boundaries of La Parguera Natural Reserve and the Boquerón State Forest are displayed.

**Figure 2:**
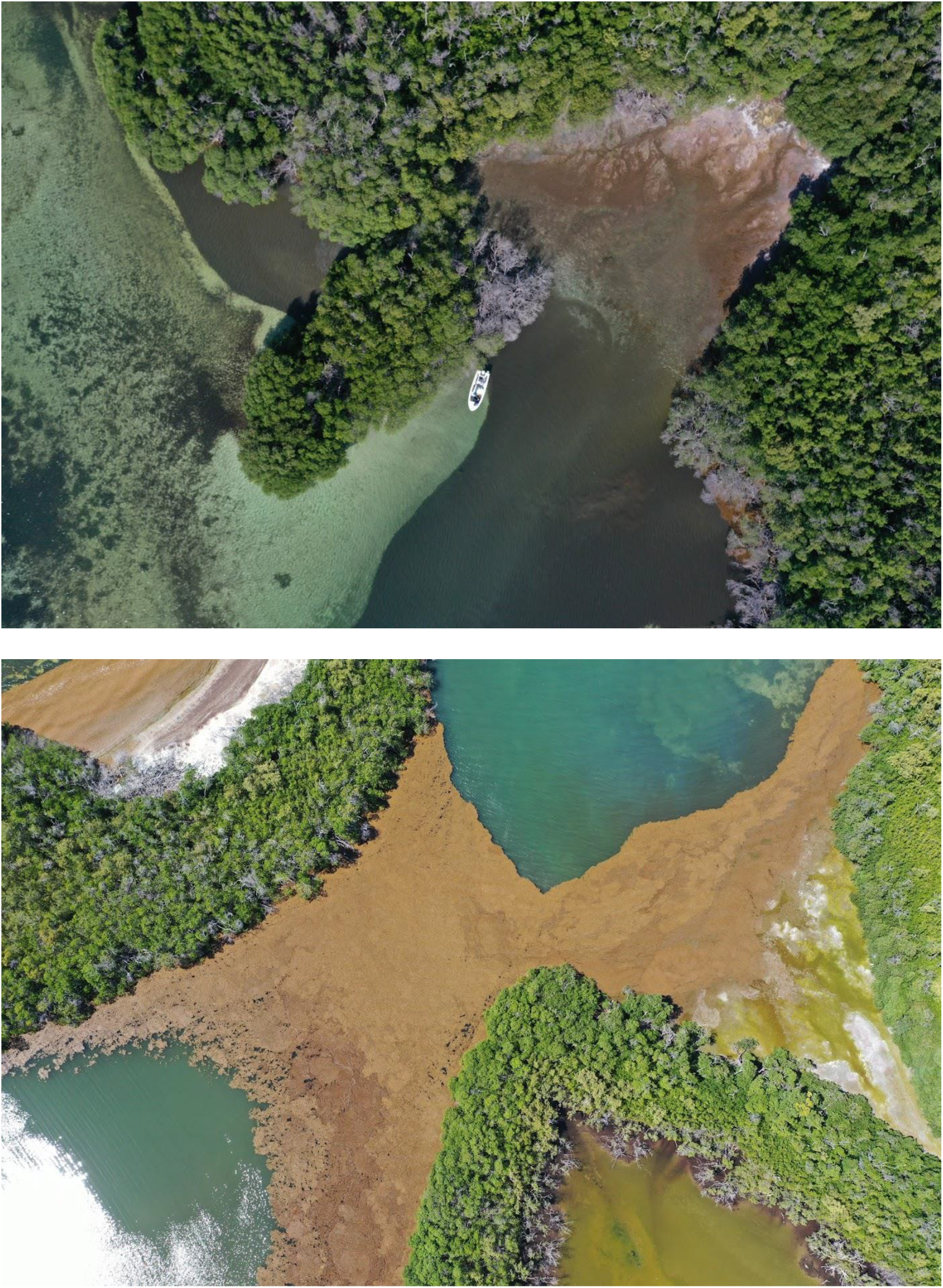
Drone image for September 2019 of Isla Guayacán, La Parguera showing *Sargassum* accumulated in the fringe mangrove areas (brown areas) and submerged in the near-shore benthic areas (black areas) (top). Drone image for July 2019 of *Sargassum* accumulated at various decomposition stages in the Isla Cueva area in fringing mangroves where this accumulation occurs (bottom). Photos by WJH.

## Materials and Methods

A very high-resolution (VHR) benthic composition and coastal vegetation map, including fringing mangroves, at the study site was developed from satellite imagery to evaluate the changes in benthic composition in La Parguera Marine Reserve potentially attributable to *Sargassum* influx and accumulations in cays, bays, inlets and near-shore environments. A previous benthic composition map was used as the baseline map for La Parguera [6]. This baseline benthic map was developed from VHR satellite imagery and field observations before the *Sargassum* influx that started in 2011.

A VHR satellite image was obtained for 2020 for the selected sites to evaluate the effect of *Sargassum* accumulation on benthic composition and coastal vegetation. To minimize the potential effects of the 2017 hurricane season on the analysis, an additional VHR image (early 2018) was also acquired. The satellite imagery selection was based on the availability of cloud-free data, timeline of the observation (*e.g.* after *Sargassum* season, after hurricane impacts), and spatial/spectral resolution of the sensors. The imagery selected and their attributes are described in Table 1.

**Table 1:**
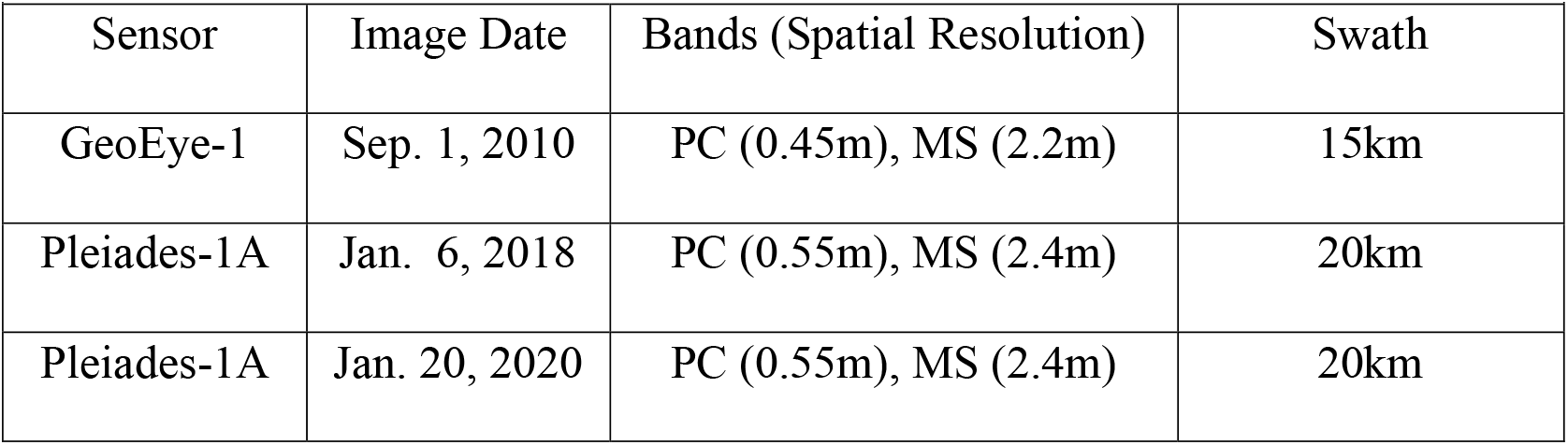
VHR Imagery Specifications. PC (Panchromatic), MS (Multi-spectral)

Imagery was pre-processed and ancillary data such as, LiDAR data (bathymetry), drone photo survey, and field observations were used to improve the classification and quantification of changes. These drone surveys were collected for various dates during the 2018 and 2019 *Sargassum* season (Figure 2). The maps produced provided for assessing changes to evaluate and quantify potential changes in the benthic composition from 2010 to 2019 and a summary of total area change in coverage for the benthic classes and coastal vegetation.

### Pre-Processing

The images were corrected for radiometric and atmospheric effects and were also pan-sharpened to improve the spatial resolution of the multispectral bands. Land and water masks were developed and applied to the images to separate the effects on coastal vegetation or benthic cover, respectively and provide and independent analysis of the changes.

The Geoeye-1 imagery was acquired from National Oceanographic and Atmospheric Administration (NOAA), National Centers for Coastal Ocean Science (NCCOS) as part of the project: Benthic Habitat Mapping off Southwest Puerto Rico (https://coastalscience.noaa.gov/project/benthic-habitat-mapping-off-southwest-puerto-rico/). A pan-sharpened orthorectified image tiles were downloaded for the selected area. Additional processing included a Dark Object Subtraction atmospheric correction [11] creating a mosaic of the tiles and a subset of the area of interest (Figure 3).

**Figure 3:**
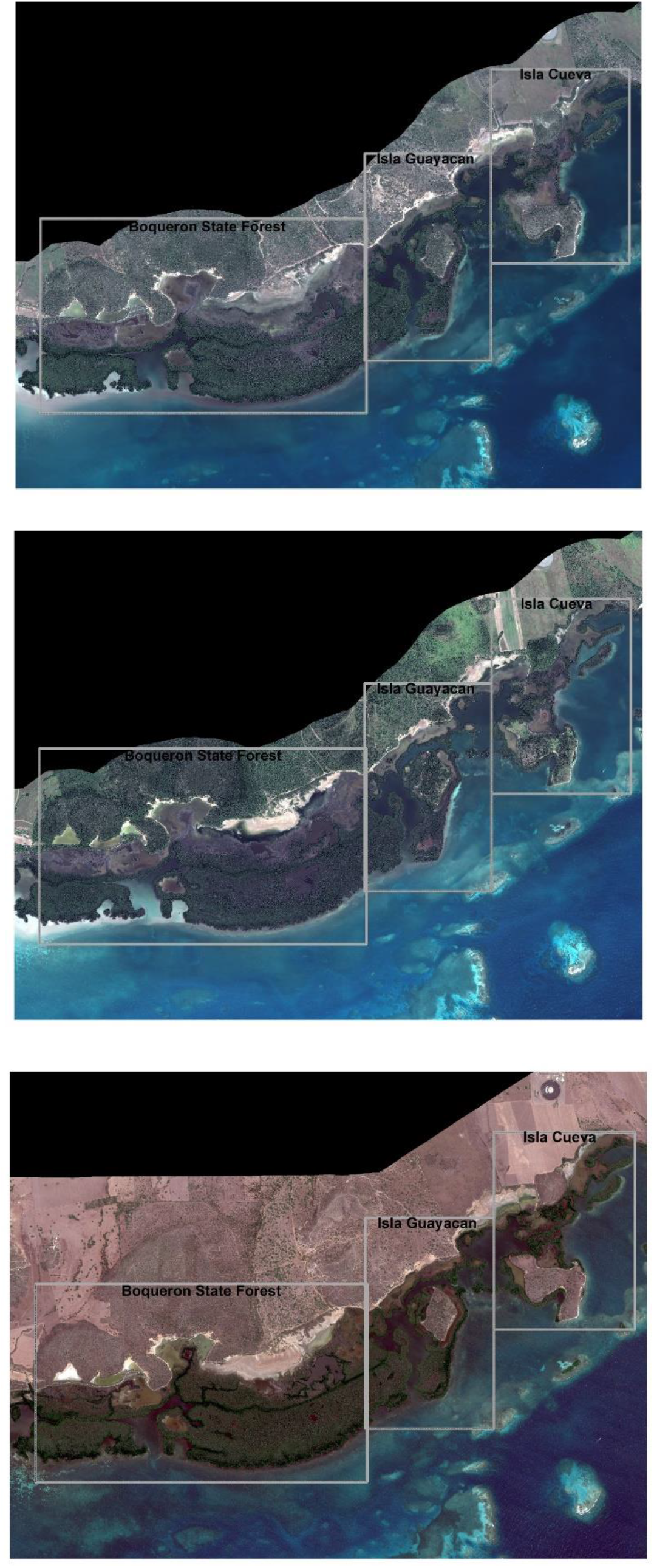
Processed Pleaides-1 imagery for 2020 (top), 2018 (middle), and the GeoEye-1 image from 2010 (bottom). The selected study areas are shown.

The Pleiades-1 images were acquired, radiometrically and atmospherically corrected using Dark Object Subtraction atmospheric correction [11] in ENVI 5.3 image processing software. A Gram-Schmidt pan-sharpening was applied to the imagery by including the panchromatic band with ground control points. Processing was completed for the January 06, 2018 and January 19, 2020 imagery to obtain a multi-spectral image at 0.55m spatial resolution (Figure 3).

#### Georeferencing

The processed GeoEye-1 image was used as the base image for georeferencing. The base image orthorectification was performed using ENVI’s Orthorectify GeoEye-1 with Ground Control module. Due to the difference in spatial resolution, the GeoEye-1 imagery was resampled to the Pleaides-1 spatial resolution of 0.55m. A subset of the Pleaides-1 imagery was developed based on the extent of the GeoEye-1 image.

#### Change Detection for Vegetation

Various techniques were used to evaluate changes in the imagery over time. For the coastal vegetation, changes in the NDVI index was used to quantify these changes. The NDVI was used here to assess the state of live green vegetation and it also indicates a level of photosynthetic activity [12]. The NDVI is calculated as follows:

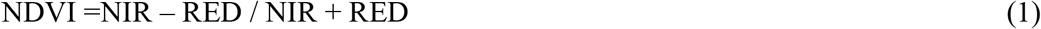

where RED and NIR are the spectral channels (bands) for the sensors in the red (visible) and near-infrared (NIR) regions, respectively. For both the Pleaides-1 and GeoEye-1 images, these correspond to the third and fourth bands. In general, the range of NDVI values are between −1.0 and +1.0, where negative values can represent water. A water mask was applied to the reduce the effects of water on the NDVI index and classification. Once the vegetation index has been processed, the new images can be subtracted from the baseline image using map algebra to evaluate the changes [7]. This was done for the 2018 and 2020 images. The NDVI pixel values were then reclassified as positive change or negative change categories if pixel values were higher (positive) or lower (negative) than the baseline image. Once the NDVI changes from the baseline were calculated, a difference image was processed between 2018 and 2020 images to evaluate NDVI changes between these dates. To calculate the net percent change of NDVI that can be attributed to the potential hurricane impacts (2018 image), or the potential Sargassum impacts (2020 image), the difference between 2010-2018- and 2010-2020-pixel count was calculated for each study area.

#### Change Detection for Benthic Composition

For the benthic composition changes, an unsupervised classification (ISODATA) as an initial evaluation, and an object-based segmentation and further classification was used to quantify the changes using all bands. A land mask was applied to the image to reduce the effects of land features on the classification. The classes and iterations were adjusted until the segmentation of benthic features was achieved. Areas with sunglint and land features still visible were grouped with the mask class. A preliminary map was obtained where major classes and major cover were classified based on the benthic classes and cover from baseline benthic cover map [6]. This initial classification step and draft benthic composition maps were produced for both 2018 and 2020 images. These preliminary benthic maps were then assigned benthic classes based on; 1) baseline survey from previous benthic map, 2) updates from boat surveys and drone images and videos for selected sites. No contextual editing [4] was applied to the segmented benthic composition polygons since the focus of the project was to quantify changes from the baseline image. Final maps classes were compared to evaluate changes in the total area by major classes and coverage from the baseline benthic maps with special interest in the potential changes in benthic cover by hurricanes (2018) and *Sargassum* (2020).

## Results

Coastal vegetation and benthic composition maps were developed to quantify changes in the study area potentially attributable to Sargassum influx and accumulations in cays, bays, inlets and near-shore environments.

### Changes in Coastal Vegetation

The processed images were evaluated for change detection using NDVI products. These products calculate the difference in NDVI from the baseline imagery (2010), to the potential hurricane (2018 image) and *Sargassum* impacts (2020 image). The total pixel area values were summarized with negative and positive change for each study area (Figure 4). Positive changes can represent an increase in NDVI pixel values due to both a “greener” vegetation (with the same coverage) or an increase of coverage (with the same greenness) form the baseline image, while negative changes can represent a decrease in NDVI pixel values due to loss of vegetation (*e.g.* canopy structure) or decrease of vegetation coverage (*e.g.* change from covered areas to barren or open water areas) [13]. Results show a dramatic increase in the positive changes in total pixel area, especially for the Boquerón State Forest (BF) from 2010-2018. However, this increase is reduced when compared to the 2010-2020 total pixel area count and can be attributed to an increase in the negative change pixel area for that period. NDVI negative changes pixel area also increased from 2010-2018 to 2010-2020 for Isla Guayacán (IG) and Isla Cueva (IC).

**Figure 4:**
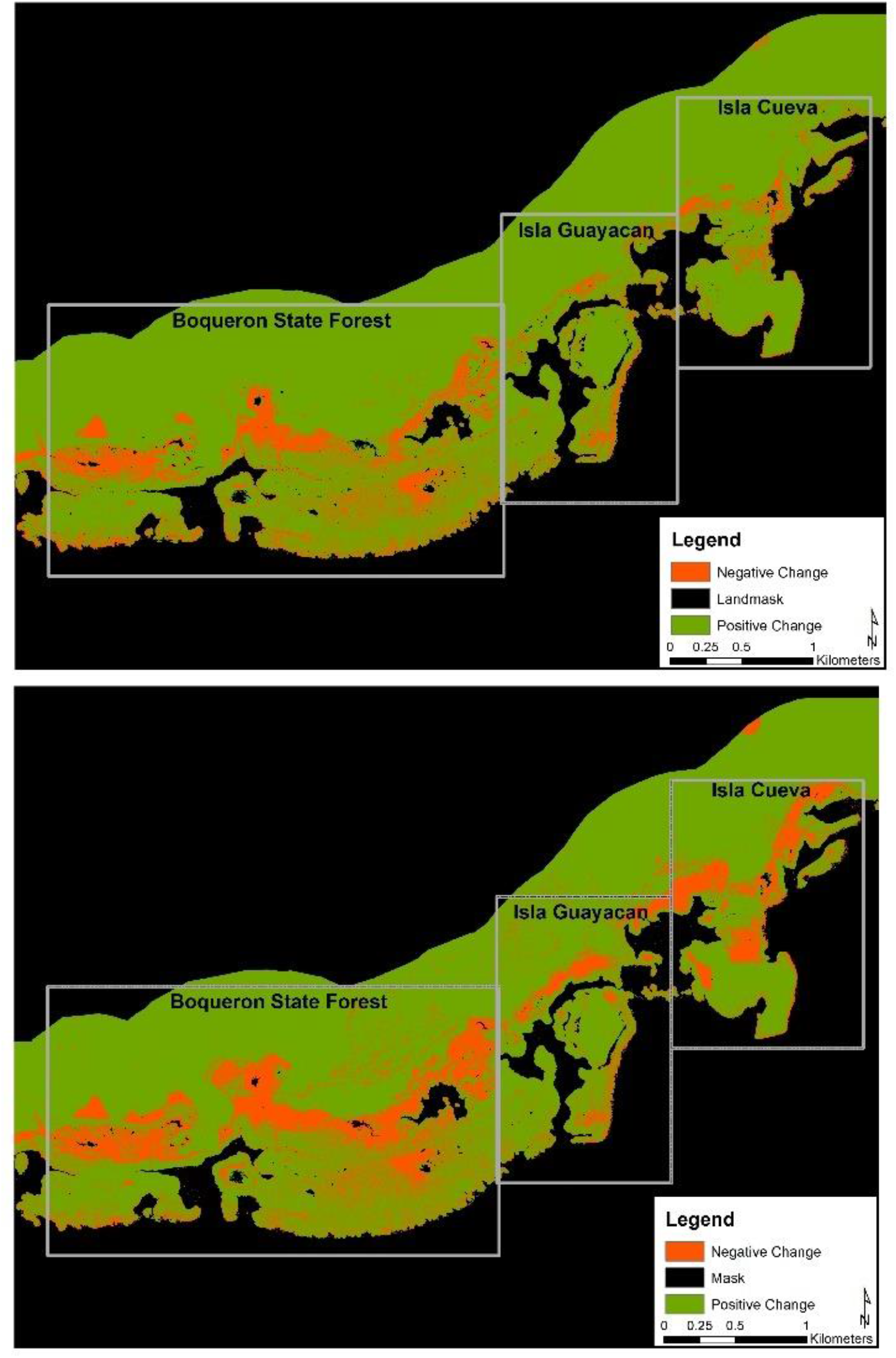
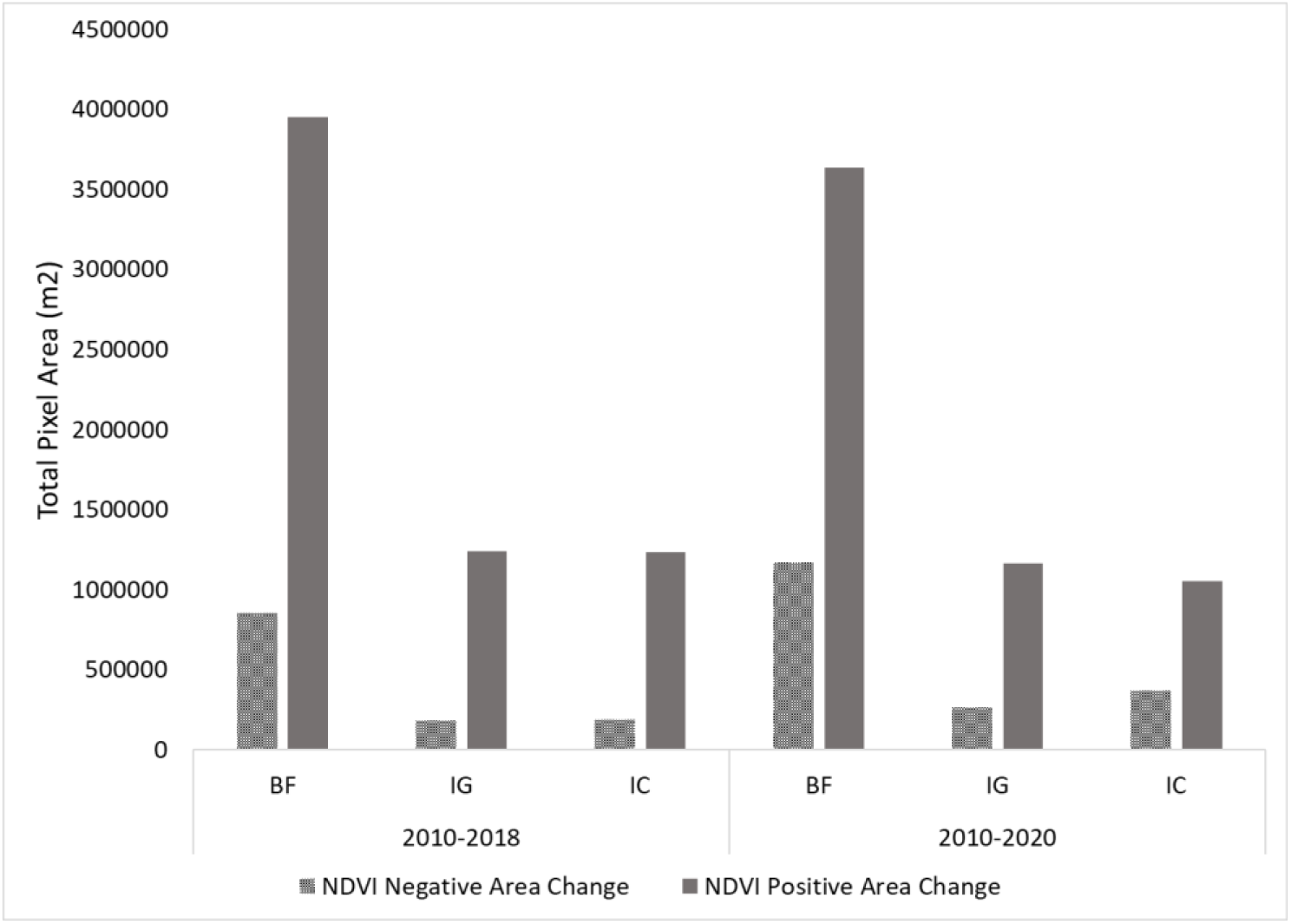
Classified NDVI imagery products for negative/positive changes for 2010-2018 (top) and 2010-2020 (middle) for each study area. A total pixel area (m^2^) summary for both NDVI products imagery for 2010-2018 and 2010-2020 (bottom).

The net % change of NDVI was calculated for each study area that can be attributed to the potential hurricane (2018 image) or *Sargassum* impacts (2020 image) (Figure 5).

**Figure 5:**
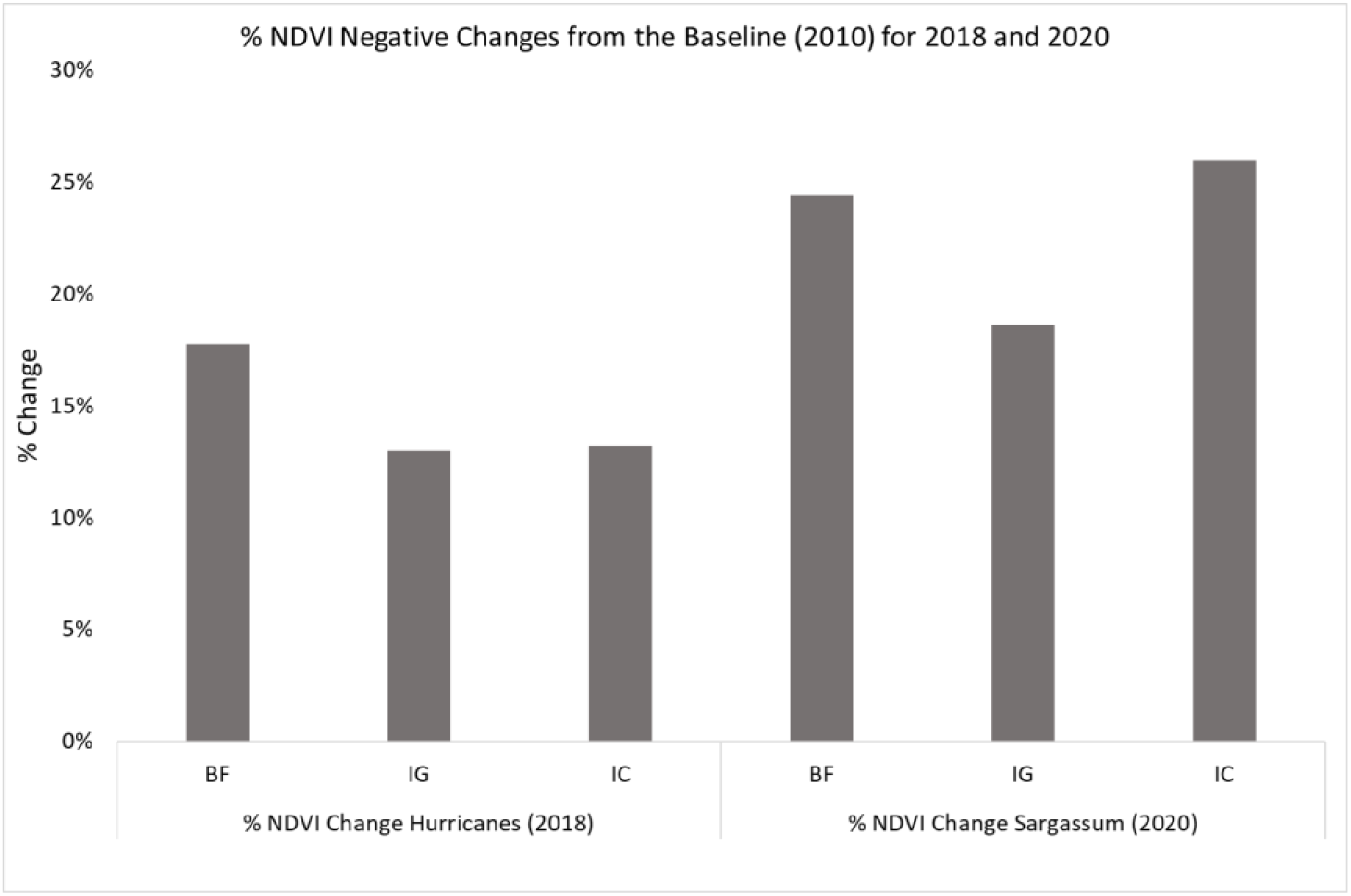
NDVI negative changes (%) from the baseline (2010) of the potential hurricane impacts (2018 image), or the potential *Sargassum* impacts (2020 image) for the selected sites.

The results show an increase in % NDVI change in the negative values from 2020 when compared with 2018. The increase in percent of NDVI negative values from 2018 to 2020 was 38% for BF, 44 % for IG, and 97% for IC.

### Changes in Benthic Composition

Final benthic composition maps were produced for both 2018 (Figure 6) and 2020 images (Figure 7), while 2010 image and NOAA benthic map are included for reference (Figure 8).

**Figure 6:**
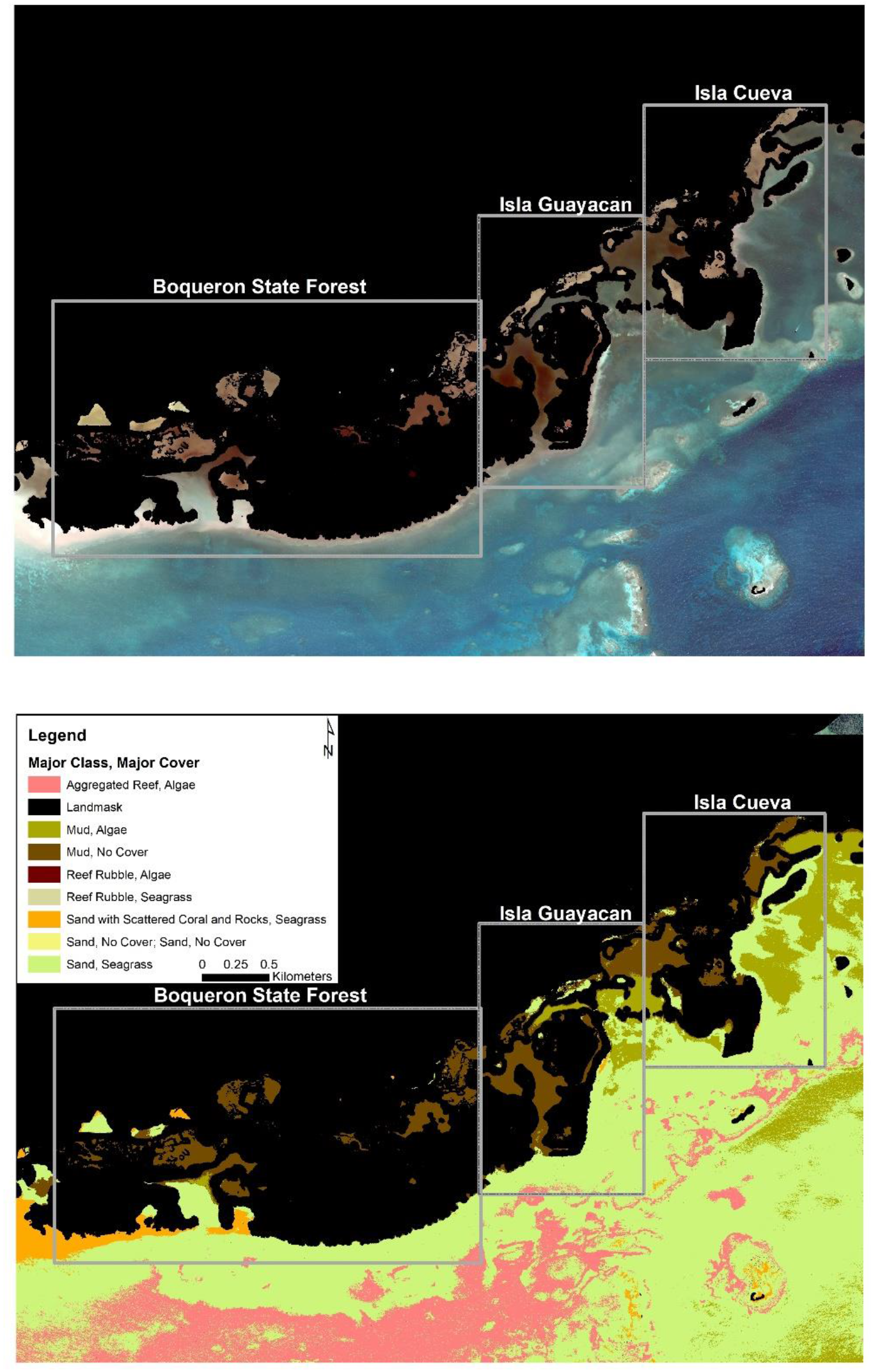
Pleaides-1 2018 true color image with landmask (top). A final classified benthic composition for 2018 showing the different benthic classes (bottom).

**Figure 7:**
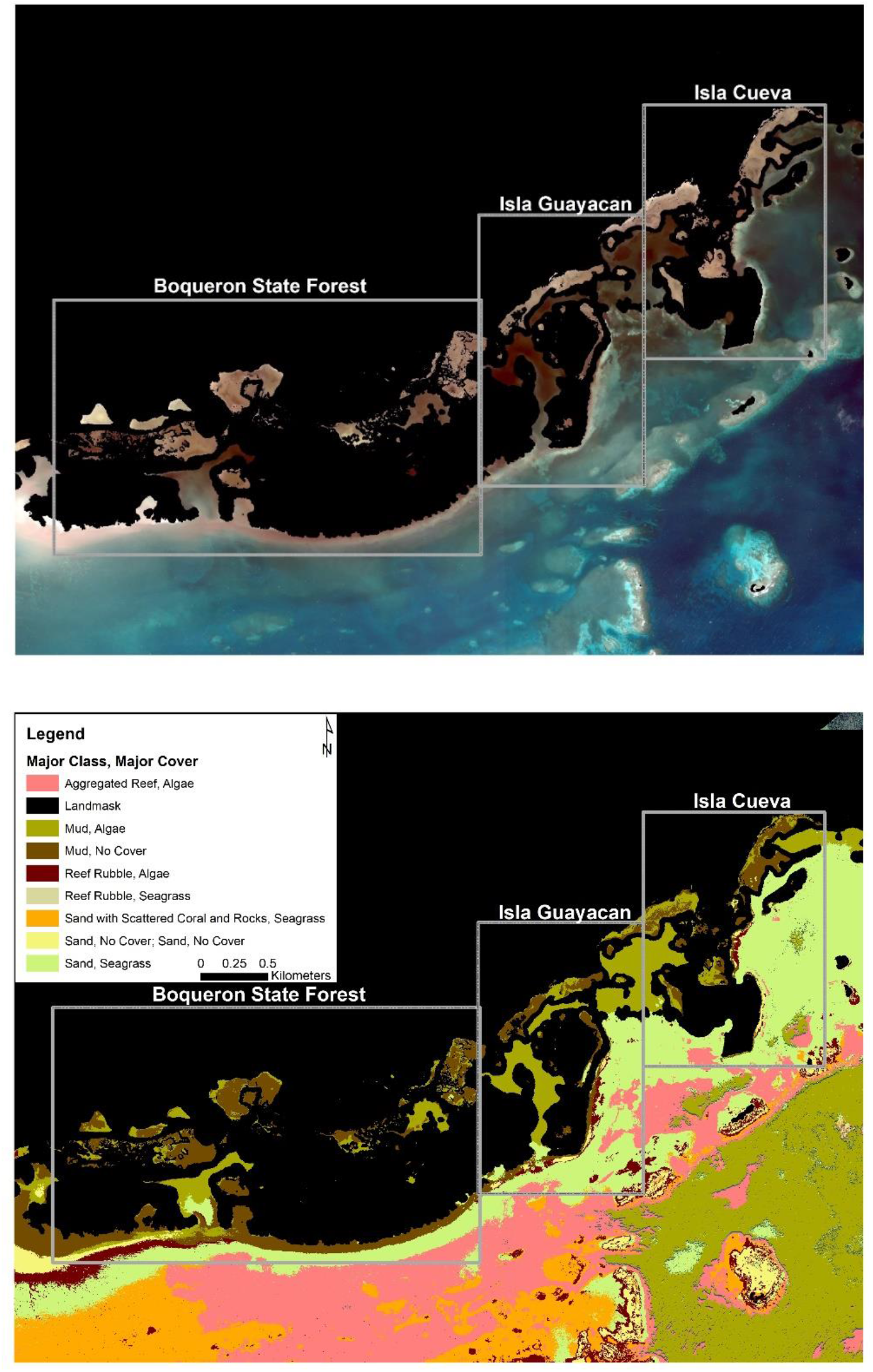
Pleaides-1 2020 true color image with landmask (top). A final classified benthic composition for 2020 showing the different benthic classes (bottom).

**Figure 8:**
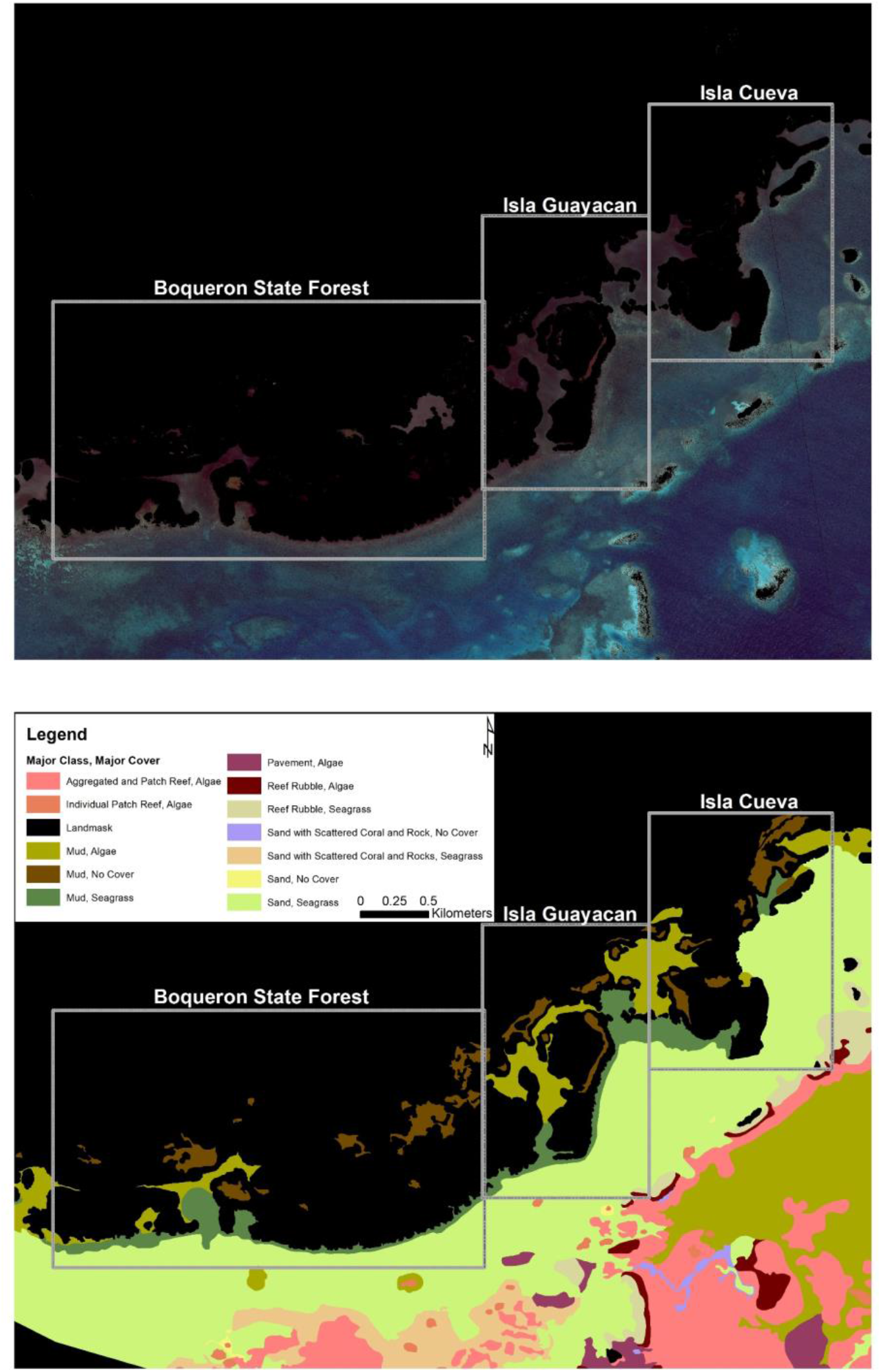
GeoEye-1 2010 true color image with landmask (top) and the final benthic composition map for 2010 showing the different benthic classes (bottom) from Bauer *et al.*, (2011).

The benthic classes were compared for 2018 and 2020 using the following categories: for the Major Classes: Aggregated Reef, Mud, Reef Rubble, Sand, and Sand with Scattered Coral and Rocks, and for the Major Cover: Algae, No Cover, and Seagrass (Figure 9). The major changes occurred as a reduction in the Sand class (12.29%), while the Mud class increased (7.37%) from 2018 to 2020. An increase in Seagrass cover (14.19%) and a reduction in the Mud cover (15.18%) was obtained from 2018 to 2020. The major cover class was compared to the baseline benthic map 2010 (Figure 10).

**Figure 9:**
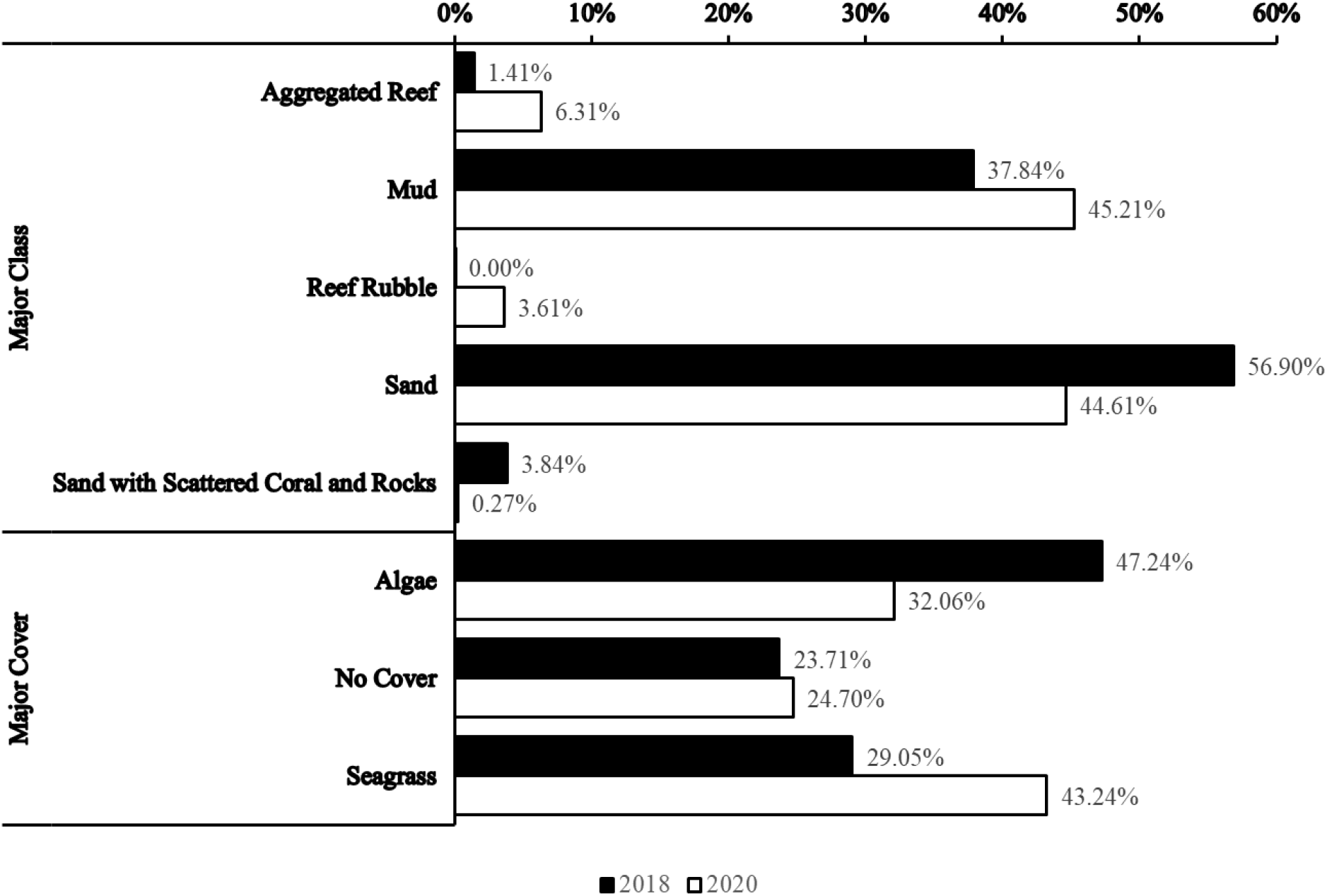
Percent of area of coverage by Major Class for 2018 and 2020, and for Major Coverage for 2018 and for 2020.

**Figure 10:**
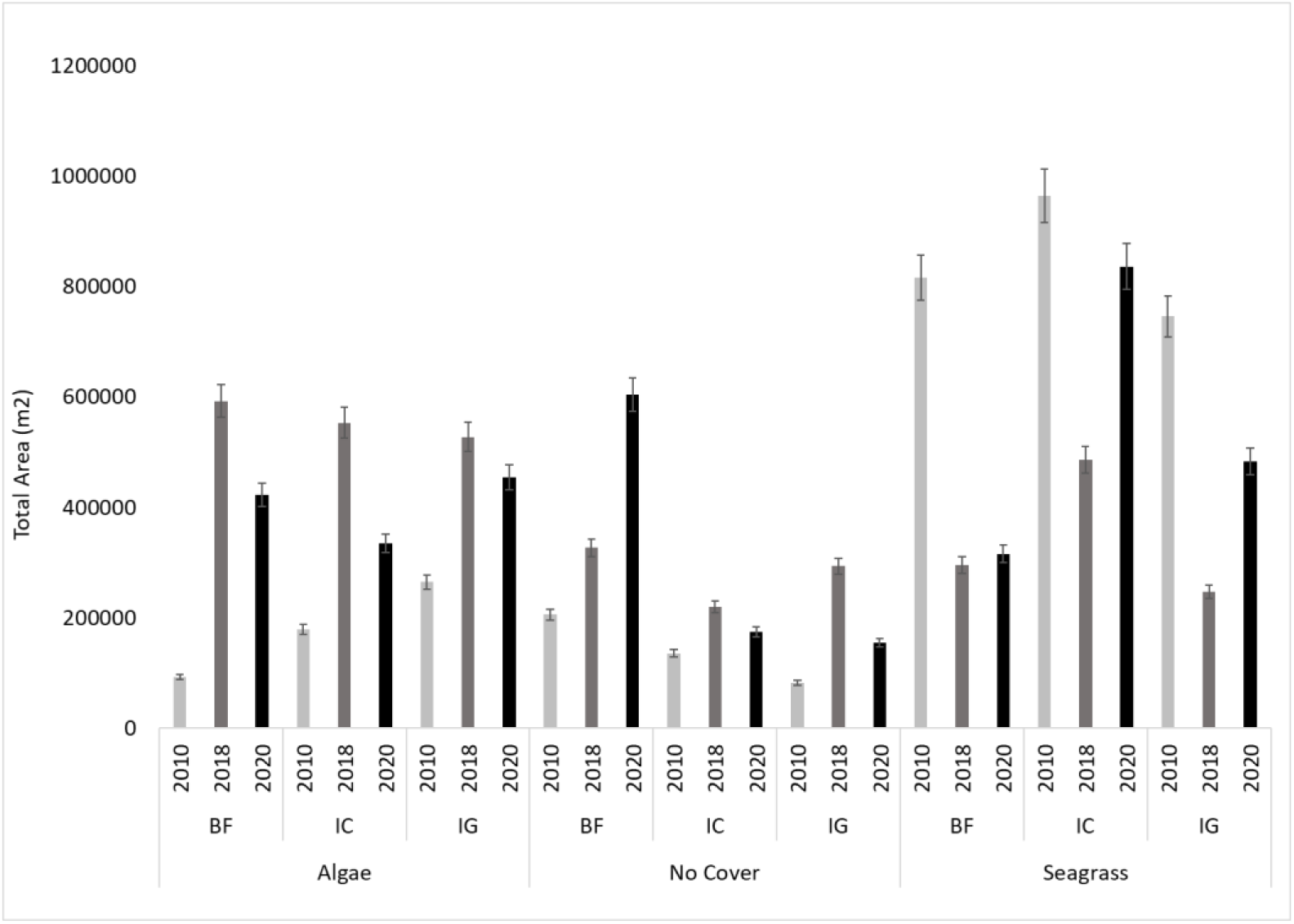
Total area of Major Cover from 2010, 2018, and 2020 for the selected study sites.

## Discussion

The coastal vegetation shows a negative trend in NDVI from 2010 to 2020 and these changes were also observed in the corresponding true color images. The major changes to the benthic composition occurred to the Sand and Mud categories, while the Major Cover category changed mainly between Seagrass and Algae.

### Changes in Coastal Vegetation

The NDVI index analysis shows a negative trend from 2010 to 2020. The total pixel area that showed a negative change in values was 546,446 m^2^, or 24% of the total area for the sites from 2010 to 2020. These changes can also be observed in the true color images from the same time period (Figure 3). Changes in the NDVI negative values from 2018 to 2020 were higher, especially for the Isla Cueva site (97%). This is consistent with the field observations and drone surveys conducted since 2018 in the area (Figure 2) that show a decrease in the standing mangrove and canopy cover extent. The NDVI index uses the Red/NIR ratios for which standing healthy vegetation produce higher values, a reduction in these values show a decrease in standing healthy vegetation or transition to barren or open areas [12]. The 2018 *Sargassum* season showed the highest biomass of *Sargassum* in the Caribbean and Intra-America Seas [1], with high accumulation occurring in the southwest coast of Puerto Rico. Floating *Sargassum* mats are driven by the southeast winds an accumulate and persist for extended periods of time, especially in Isla Cueva and Isla Guayacán sites due to their geographic orientation. Also, a substantial influx of *Sargassum* can extend deep inside the fringing mangrove forest during high tide that persists and decomposes there, which can lead to anoxic conditions with potential changes to the biochemistry and hydrodynamics of the mangrove root system [3].

Hurricane effects could also have impacted the NDVI values, since the percentage of negative values was higher than the baseline (2010), especially for the Boquerón Forest site. The authors evaluated the NDVI values for Puerto Rico after Hurricane Maria (September 2017) and showed that NDVI values recovered to pre-hurricane levels around 1.5 months (November 2017) after the hurricane impacts, and that the vegetation farthest from the storm center and with lower elevation were the least affected [7]. This information is confirmed by the authors [14] that shows that mangroves were severely impacted from Hurricane Maria in the eastern part of the island when compared to other areas. Also, canopy coverage from mangrove forest recover to 60% pre-hurricane conditions between 3–6 months post-storm for Hurricane Irma impacts in Florida [15]. The image used for our NDVI analysis was from January 2018 which suggest that NDVI values and canopy cover were back to pre-hurricane levels, and that negative NDVI are only showing the areas severely impacted by the hurricane.

Several limitations can be found when using NDVI for change detection analysis that include atmospheric effects, new leaf growth in crops, and seasonality [12]. However, these limitations were minimized in the image pre-processing. A water mask was applied to the images to remove any potential NDVI negative values from water to the pixels. In addition, analysis of changes was completed from the baseline image to ensure a standardization of the differences for the 2018 and 2020 images. In addition, the 2018 and 2020 images were selected from the same period (January) to remove any differences in seasonality that might affect the “greenness values” that are obtained from the NDVI.

### Changes in Benthic Composition

Benthic composition maps were developed for 2018 and 2020 imagery to evaluate potential impacts to the benthic community structure from both hurricanes and *Sargassum*. The main changes to the Major Class category occurred to the Sand and Mud categories, while the Major Cover category changed mainly between Seagrass and Algae. No major changes were observed to the No Cover class from 2018 to 2020 (Figure 10). The major changes from 2018 and 2020 occurred mainly in unconsolidated sediments (*e.g.* Sand, Mud) and submerged aquatic vegetation (*e.g.* seagrass, algae), both of which can have similar spectra and can be very difficult to differentiate from multi-spectral imagery [16]. Due to these considerations, the changes from 2018 and 2020 cannot be considered relevant since the No Cover class remained relatively similar for the time period. Benthic structure impacts from storm surge of hurricane Maria and Irma should have been minimum, due to the storm track and distance from study area [7]. Also, the Puerto Rico central mountain chain can create a shade effect, that may have provided additional protection from the storm surge at out sites [17].

Using the 2018 and 2020 true color VHR imagery combined with the drone observations, we could locate the areas where the *Sargassum* accumulated, decomposed, and deposited (Figure 11). These selected areas at Isla Cueva and Isla Guayacán show the changes in accumulation of *Sargassum* from 2018 to 2020, which can also be identified in the drone imagery from September 2019. The accumulation of *Sargassum* persisted and was visible in the 2020 satellite imagery even after the end of the 2019 *Sargassum* season.

**Figure 11:**
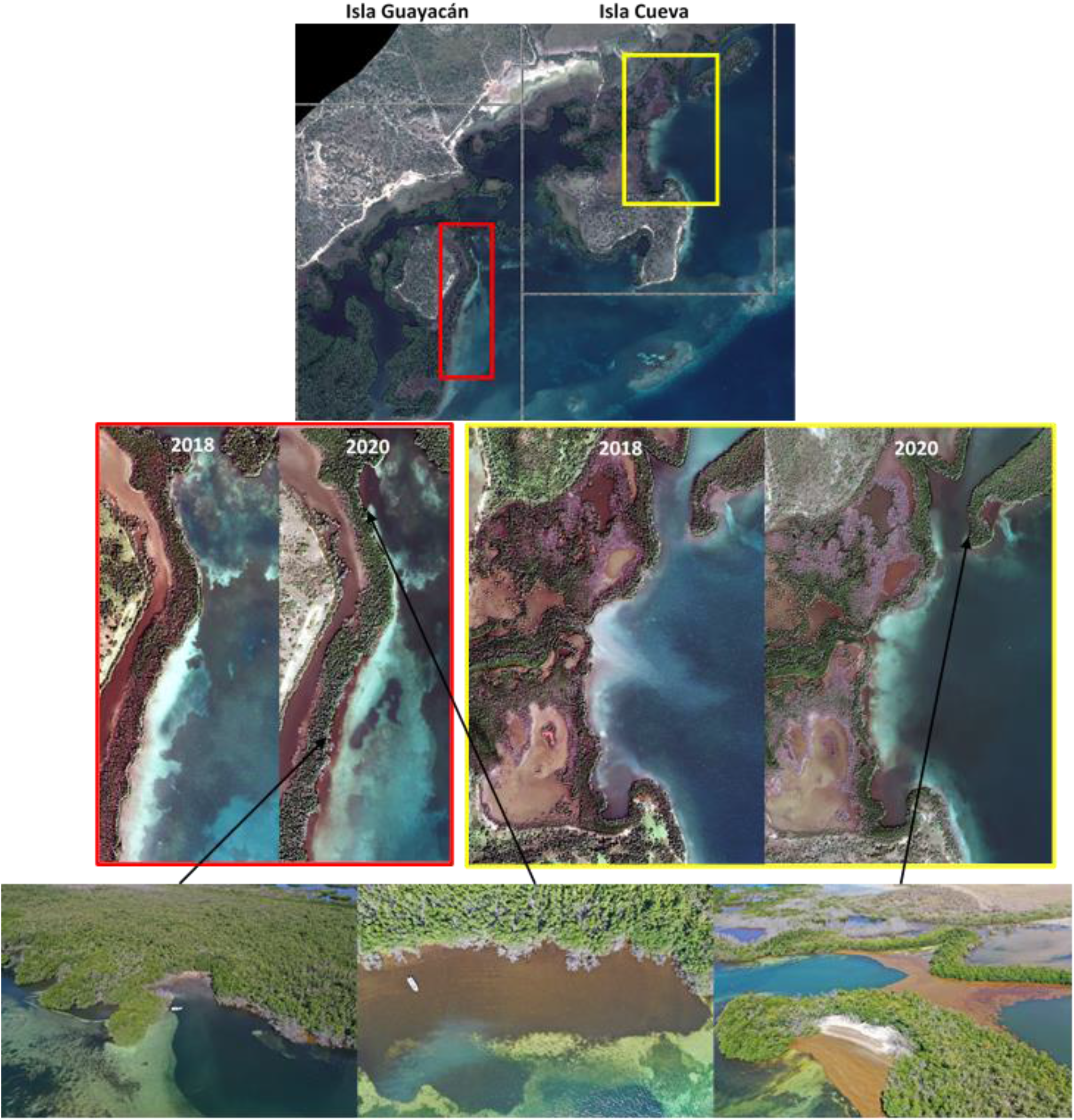
Enhanced areas from satellite imagery (top) from Isla Guayacán in red box (middle-left) and Isla Cueva in yellow box (middle-right) for both 2018 and 2020 showing changes in accumulated *Sargassum* at various locations. Drone images from September 2019 from Isla Guayacán (bottom-left-middle) and Isla Cueva (bottom-right) location on satellite imagery showing the accumulations of *Sargassum*.

The decomposed *Sargassum* in the benthos was not captured by the benthic classification, probably due to the lack of differentiation from unconsolidated sediments like mud. Quantifying the benthic changes presented some additional challenges due to the water column effect on the signal received by the sensors. Additional factors such as sunglint and sediment resuspension, especially in the 2018 image may have limited the classification process.

Our study found major changes to the coastal ecosystems of La Parguera (*e.g.* fringing mangroves and seagrass) which it’s interdependence results in a highly productive and biodiverse marine resource [18]. Hurricane impacts to coastal vegetation and benthic ecosystems have been documented [15,7], but these can still be considered episodic events providing time for the ecosystem to recover. However, the cumulative and chronic impacts of *Sargassum* [3] may not provide a recovery time to these coastal ecosystems, especially if the system has been severely impacted by a hurricane or storm. The changes to the coastal vegetation and benthic composition we observed in our study occurred in a short time frame, considering that both Sargassum [1] and extreme weather events such as storms and hurricanes [19, 20] will continue and potentially increase, natural resource managers need to consider the combination of these scenarios and its potential long-term impacts to the ecosystems.

## Conclusions

Significant changes in coastal vegetation from 2010 to 2020 were quantified in the study sites in La Parguera area using the NDVI and it was possible to differentiate between the hurricane and *Sargassum* impacts. Further studies should focus on how these accumulations may have a detrimental effect in the biochemistry and hydrodynamics of the mangrove root system, which may result in loss of mangrove forest habitat. Although we quantified that the observed percent increase in the NDVI negative values could be due to *Sargassum*, other factors like hydrological changes to the mangrove forests may have also contributed to these changes. Debris accumulations, sediments redistribution or wind damage to canopy structure [14,15] can have a cumulative effect, which cannot be resolved with the temporal scales used in this study.

Benthic composition maps were completed for the 2018 and 2020 imagery and were used to evaluate the potential changes from 2010 to 2020. However, similarities in the spectral characteristics of the substrate limited the reliability of the retrievals and could not differentiate the accumulated *Sargassum* in the benthos. Drone surveys using multi-spectral cameras combined with optical field surveys may provide a more detailed optical information and the high-resolution and temporal scale needed.

This approach provides a quantifiable method to evaluate *Sargassum* impacts to the coastal vegetation and benthic composition using change detection of VHR images, and to separate these effects from other extreme events.

## Acknowledgements

The authors would like to thank Mariana León-Pérez for reviewing and providing recommendations to the manuscript. This study was supported in part by the Caribbean Coastal Integrated Ocean Observing System (CARICOOS) award number NOAA NA16NOS0120026. This study was supported in part by The National Oceanic and Atmospheric Administration ‒Cooperative Science Center for Earth System Sciences and Remote Sensing Technologies (NOAA-CESSRST) under the Cooperative Agreement Grant # NA16SEC4810008. The statements contained within the manuscript/research article are not the opinions of the funding agency or the U.S. government but reflect the author’s opinions. The funders had no role in study design, data collection and analysis, decision to publish, or preparation of the manuscript.

